# FLARIM v2.0, an improved method to quantify transcript-ribosome interactions *in vivo* in the adult *Drosophila* brain

**DOI:** 10.1101/2021.08.13.456301

**Authors:** Petra Richer, Sean D. Speese, Mary A. Logan

## Abstract

Neural injury triggers striking immune reactions from glial cells, including significant transcriptional and morphological changes, but it is unclear how these events are coordinated to mount an effective immune response. Here, we present a new variant of the <u>Fl</u>uorescence <u>a</u>ssay to detect ribosome interactions with <u>m</u>RNA (FLARIM), which we term FLARIM v2.0, to visualize single immune gene transcripts and association with ribosomes in glia responding to neurodegeneration. Specifically, using an *in vivo* axotomy assay in *Drosophila*, we show that matrix metalloproteinase-1 (Mmp-1) mRNAs and associated ribosomes are detected in distal processes of reactive glia where they are actively engulfing degenerating axonal material, suggesting that local translation is an important component of the glial immune response to axotomy. This work also validates our enhanced FLARIM assay as a promising tool to investigate mechanisms of mRNA transport and translation in a wide range of *in vitro* and *in vivo* paradigms.

## Introduction

Glia are the resident immune cells of the brain and respond swiftly to neuronal trauma, pathogenic insult, and degeneration(1-3). Following neuronal damage, activated glia undergo distinct transcriptional, morphological, and functional changes(4-6). In many cases, reactive glia are neuroprotective, releasing pro-survival factors and clearing damaged neurons through phagocytic engulfment(7, 8). Thus, understanding how glial immune responses are activated and carried out will offer new insight into approaches that could delay the onset or progression of a range of neurodegenerative disorders and injury conditions.

The fruit fly, *Drosophila melanogaster*, offers a powerful genetic *in vivo* model to explore evolutionarily conserved glia-neuron signaling events after neural injury(9-11). For example, after axotomy, adult *Drosophila* axons undergo a classic Wallerian degeneration (WD) program, and local glial cells display striking immune responses to invade injury sites and rapidly clear degenerating axonal material(12-14). Notably, nerve injury triggers robust transcriptional changes, including upregulation of the conserved glial engulfment receptor Draper and the secreted protease matrix metalloproteinase-1 (Mmp-1)(12, 15-22).

Our lab has recently shown that upregulation of Mmp-1 is necessary for timely glial clearance of degenerating axonal projections in the olfactory system of adult flies(22). Mmp-1 is secreted from local ensheathing glial cells, likely to facilitate extracellular matrix remodeling, allowing glia to extend membrane processes into neuropil regions and clear degenerating olfactory receptor neuron axons(22, 23). Notably, the cell bodies of ensheathing glial cells responding to degenerating axons in the olfactory system do not enter neuropil regions, which raises interesting questions about how key immune molecules are released within the neuropil at injury sites(12, 24). Previous work in other model systems has indeed demonstrated that directed mRNA transport and local translation are important for glia to carry out normal functions. For example, in oligodendrocytes, myelin basic protein transcripts are localized to oligodendrocyte processes to adequately myelinate axons in an activity-dependent manner, while astrocytes have been shown to influence interactions at tripartite synapses and the gliovascular interface through a subset of discrete localized and locally translated transcripts(25-29).

In order to further explore the transcriptional and translational changes that are essential for proper glial responses to damaged axons, our lab has utilized various single molecule fluorescence *in situ* hybridization (smFISH) techniques for the detection of individual transcripts and a new variation of <u>Fl</u>uorescence <u>a</u>ssay to detect ribosome interactions with <u>m</u>RNA (FLARIM), referred to as FLARIM v2.0, to detect ribosome association with *mmp-1* transcripts in glial cells following nerve injury(30). smFISH has been employed in glial cells and various *Drosophila* tissues to detect individual mRNAs; however, previous studies have not yet employed mRNA and ribosome detection together in the context of neuronal injury *in vivo*(31-38). Here, we have utilized single molecule inexpensive FISH (smiFISH), hybridization chain reaction (HCR), and FLARIM, in order to visualize *mmp-1* transcript localization and translation in reactive glia following axotomy(30, 39-41). Our findings reveal that *mmp-1* transcripts are localized to and associate with ribosomes at distal glial processes at injury sites, suggesting that local translation of Mmp-1 may be an important mechanism by which glia access and phagocytically clear neuronal debris.

## Results

### Ensheathing glia respond to olfactory receptor neuron injury in *Drosophila*

Axotomy in the adult *Drosophila* antennal system is a well-characterized model to investigate the molecular and cellular underpinnings of axon degeneration and glial immune responses (Fig. 1A)(12, 42). The olfactory receptor neuron (ORN) cell bodies reside in the external olfactory organs of the fly, the antennae and maxillary palps, and project to the antennal lobes of the central brain, where they synapse onto second order neurons(43, 44). Surgical removal of the antennae and maxillary palps results in axotomy of the ORNs, which undergo typical WD over the course of days, after which they are cleared by the surrounding glial cells(12, 45). In the antennal lobes, glial cell bodies (ensheathing glia and astrocytes) are located at the very edges of the antennal lobes and extend projections to closely associate with neuropil regions (axons, dendrites, and synapses) and demarcate olfactory glomeruli (Fig. 1B)(9, 46). In context of ORN injury, ensheathing glia have been demonstrated to phagocytose and clear degenerating projections(24).

**Figure 1:**
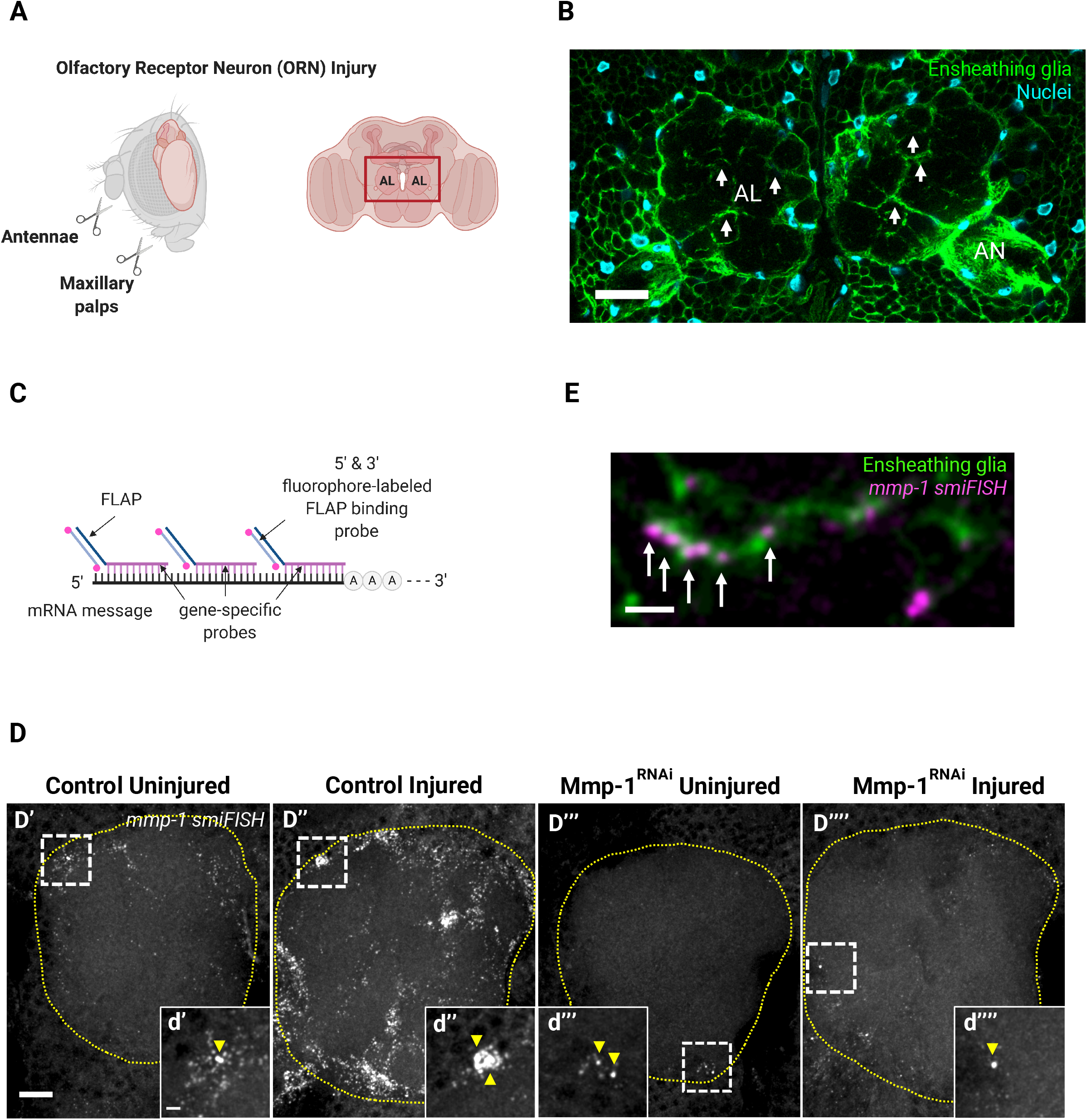
*mmp-1* transcripts are upregulated in ensheathing glia following neuronal injury. **(A)** Schematic representation of the olfactory receptor neuron (ORN) injury, demonstrating the removal of the 3^rd^ antennal segments and maxillary palps. ORN cell bodies in the aforementioned structures extend their projections to the antennal lobes (ALs) of the central fly brain (red box). Created with BioRender.com **(B)** Representative image of ensheathing glia surrounding the ALs in an uninjured brain. White arrows indicate ensheathing glial fine processes within ALs. AL: antennal lobe, AN: antennal nerve. Single slice (1µm). Scale bar: 20µm. Genotype: *;UAS-mCD8::GFP/UAS-mCD8::GFP;TIFR-Gal4/UAS-LacZ::NLS*. **(C)** smiFISH schematic. Created with BioRender.com **(D’-D’’’’)** Representative images of *mmp-1* smiFISH in uninjured and injured (4.5hpi) single ALs (yellow dotted line). Images are MIPs: 3.75µm. Scale bar: 10µm. Control genotype: *;tub-Gal80*^*ts*^*/+;repo-Gal4/+*. Mmp-1^RNAi^ genotype: *;tub-Gal80*^*ts*^*/UAS-Mmp-1*^*RNAi*^*;repo-Gal4/+*.**(d’-d’’’’)** High magnification images of *mmp-1* smFISH (white box), representing single transcripts and transcription start sites (yellow arrows). MIP: 3.75µm. Scale bar: 2µm. **(E)** High magnification image of *mmp-1* transcripts in ensheathing glial processes (white arrows) 4hpi. Single slice: 0.43µm. Scale bar: 2µm. Genotype: *;UAS-mCD8::GFP/UAS-mCD8::GFP;TIFR-Gal4/UAS-LacZ::NLS*.

Individual ensheathing glial cells have highly complex morphologies, with extensive fine processes that interact with multiple glomeruli within the antennal lobe (Fig. 1B)(46-48). In the event of an injury, the cell bodies of ensheathing glia remain positioned at the outskirts of the antennal lobes, while the processes further invade glomeruli to clear degenerating ORNs(12, 22, 24). This raises the question of how intracellular signaling is mediated within ensheathing glial cells from the cell body to the distal processes after an injury event. In order to explore this question, we utilized multiple single molecule FISH (smFISH) methods to study how a well characterized immune gene, *mmp-1*, is localized and regulated at the transcriptional and translational level following ORN axotomy.

### smiFISH reveals upregulation and localization of *mmp-1* transcripts following neuronal injury

Our previous RNAseq and qPCR studies have demonstrated that the *mmp-1* transcript is acutely upregulated in the central nervous system (CNS) and ventral nerve cord (VNC) of the fly following neuronal injury. As prior research has shown that *mmp* transcripts can be localized to subcellular compartments and locally translated in an activity dependent manner, we wanted to determine the localization of *mmp-1* mRNA in ensheathing glia following ORN axotomy(49, 50). In order to accomplish this, we utilized smiFISH to detect the *mmp-1* transcript following injury (Fig. 1C)(40). The Mmp-1 smiFISH probeset included 41 probes and was designed utilizing Oligostan (Supplementary Table 1)(40). Our previous RNAseq and qPCR data suggested that there is a strong transcriptional response in the ensheathing glia 3 hours post ORN axotomy(22). As a result, we used this as a starting point for our smiFISH studies.

In accordance with our qPCR and RNAseq data, we found that smiFISH reveals a low level of Mmp-1 transcript present in the uninjured brain, with the highest amount of transcript being expressed in cells surrounding the antennal lobe, likely ensheathing glia (Fig. 1D). In response to injury, we observe a significant increase in the amount of *mmp-1* transcript detected at 4.5hpi (Fig. 1D’’). Interestingly, when labeling ensheathing glial membranes with GFP, we also observed the localization of the *mmp-1* smiFISH signal to fine processes following injury (Fig. 1E).

To determine if the Mmp-1 probeset is specific to the *mmp-1* transcript, we performed an injury experiment where Mmp-1 was knocked down in all glia using RNAi. As expected, the RNAi knockdown eliminated most of the smiFISH signal (Fig. 1D’’’-D’’’’). We did however detect large spots of Mmp-1 transcript right outside the antennal lobe, where the nuclei of ensheathing glia are located (Fig. 1d’-d’’’’). We posit that these large intense puncta represent the site of Mmp-1 transcription in the nucleus, as the transcript would not be subject to RNAi mediated degradation until it was transported to the cytoplasm(40, 51).

### The FLARIM v2.0 method can be used to detect mRNAs and associated ribosomes

While the smiFISH approach allowed us to localize the *mmp-1* transcript to distal glial processes following neuronal injury, it does not give us insight into whether there may be local translation of the transcript at distal sites. Currently, the most accepted method for investigating local translation is the use of photoconvertible fluorescent timer proteins, however this approach requires tagging the protein of interest in the context of native mRNA with all its regulatory 5’ and 3’ elements(52, 53). More recently, a non-transgenic approach was developed to assess the ribosome load on a transcript of interest while maintaining spatial localization within the cell. This method, <u>Fl</u>uorescence <u>a</u>ssay to detect ribosome interactions with <u>m</u>RNA (FLARIM), utilizes pairs of oligonucleotide probes that bind separately to ribosomes and to the mRNA of interest (Fig. 2A)(30). When these probes are in close proximity they form a full sequence for a linker probe carrying a complete <u>H</u>ybridization <u>C</u>hain <u>R</u>eaction (HCR) initiator (Fig. 2A)(39, 41).

**Figure 2:**
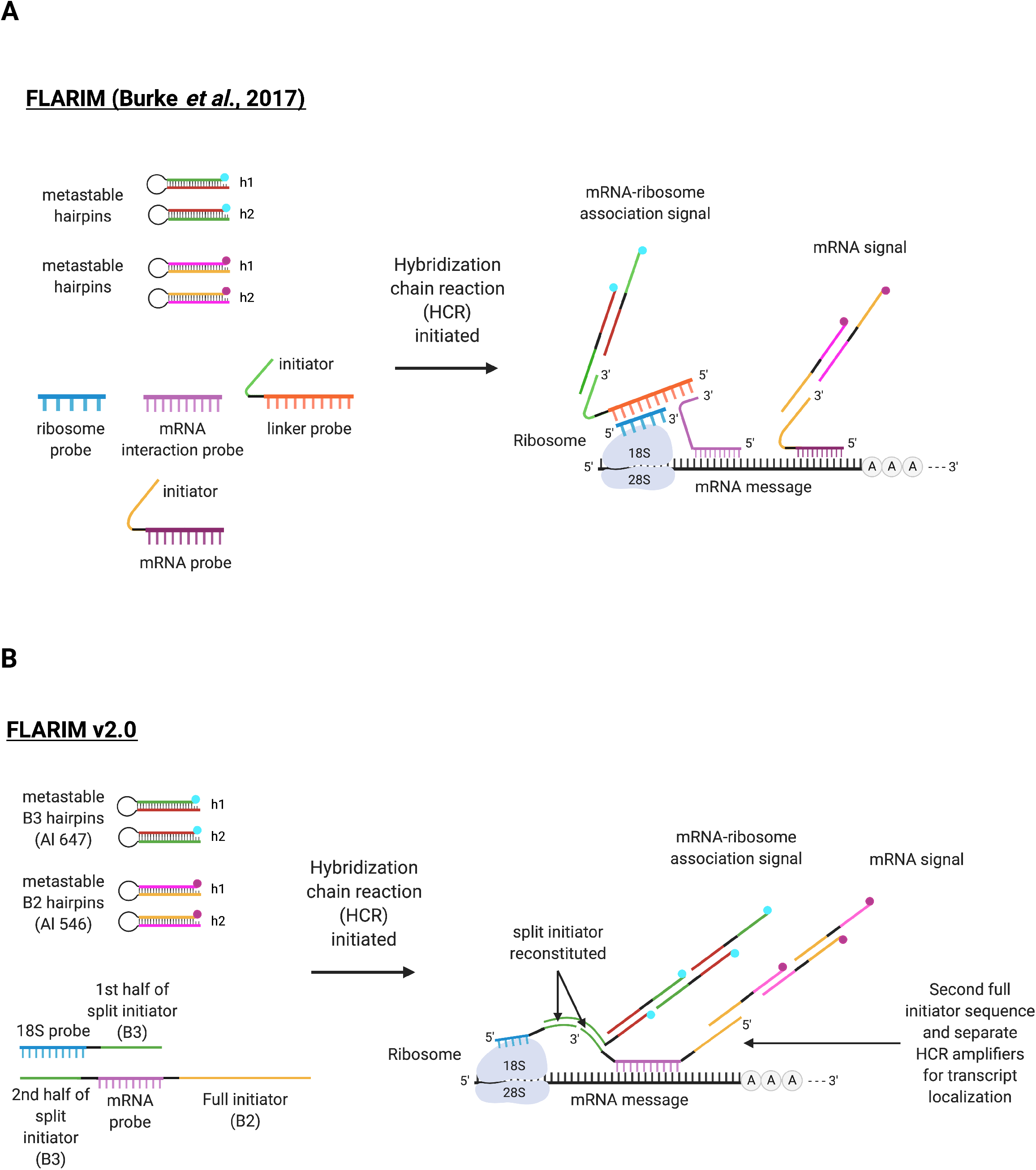
Novel FLARIM v2.0 probeset. **(A)** FLARIM (Burke *et al*., 2017): Utilizes 4 probes to achieve distinct mRNA and ribosome-specific signals by HCR amplification and allows for assessment of ribosome load on transcripts of interest. **(B)** FLARIM v2.0: Utilizes 2 probesets: 1) an 18S ribosome-specific probe with the first half of a split B3 initiator and an mRNA-specific probe with the second half of the split B3 initiator and 2) a full B2 initiator on the 5’ end of the gene specific probe. This B2 initiator allows for transcript localization and aids in quantification by creating a normalization channel for the FLARIM signal Hybridization with these probesets and amplification using metastable hairpins generates a separate fluorescent signal for mRNA and mRNA-ribosome association detection *in vivo*. Created using BioRender.com.

While the original method was able to robustly assess the ribosome load on transcripts of interest, it has never been used *in vivo* on whole mount tissue. Moreover, the use of the linker probe requires a 2^nd^ round of hybridization before initiating the HCR reaction. Lastly, the original method made use of separate pools of gene specific probes for transcript localization and ribosome association, which can be problematic on short transcripts where probe binding sites may be limiting (Fig. 2A). To address these drawbacks, we have utilized split initiator technology to include half an HCR initiator, split B3 initiator, for reporting ribosome association on the 3’ end of the gene specific probe and the other half of the B3 initiator on the 3’ end of a set of 25 probes that will hybridize to the 18S rRNA (Supplementary Tables 2 and 3)(39, 41). Finally, we tagged the gene specific probes with a different full HCR initiator (B2) on the 5’ end to allow for transcript localization (Supplementary Table 3)(39, 41). By taking advantage of new developments in oligonucleotide synthesis, Integrated DNA Technologies’ oPools Oligo Pools service, we are able to synthesize a probeset for a gene of interest for ∼$100 USD. We are calling this modified version FLARIM v2.0, and we herein demonstrate that this relatively inexpensive method is able to achieve transcript localization and ribosome association at depth in whole mount tissue (Fig. 2B).

In FLARIM v2.0, probes are first hybridized to the transcript of interest and to 18S ribosomes. Then, during the detection stage, fluorescent metastable hairpins are used to generate signals via HCR (Supplementary Fig. 1). On the gene-specific probe, the full initiator sequence opens up a set of hairpins to create a fluorescent signal indicating the presence and localization of the mRNA transcript. Moreover, when the split initiators of the 18S ribosome probes and the gene-specific probes are reconstituted, a second set of hairpins are opened, creating a distinct fluorescent signal to allow for mRNA-ribosome association detection (Fig. 2B).

The dual detection method produces diffraction limited spots, if amplification is carried out for shorter periods of time and allows for transcript and ribosome detection in a cell-type specific manner, providing spatiotemporal resolution. The fluorescent tags on hairpins can also be varied to accommodate multiplexing within the same sample. Additionally, FLARIM v2.0 can be used to investigate mRNA localization and local translation in response to various stimuli, such as neuronal injury. We decided to assess the regulation of *mmp-1* following axotomy using the FLARIM v2.0 method.

### Application of the FLARIM v2.0 method in the *Drosophila* ORN injury model

In order to validate our Mmp-1 probeset and explore how an immune gene is regulated following axotomy, the FLARIM v2.0 method was tested in the context of the ORN injury model (Fig. 3A). In uninjured brains, transcripts are sparse and localized to the edges of the antennal lobes, where ensheathing glial cell bodies are located. Some transcripts are also associated with ribosomes, generating a FLARIM signal and indicating that some translation may be occurring under basal conditions. In brains that have undergone an ORN injury, there is a robust upregulation of the *mmp-1* transcript (magenta) and ribosome-association (cyan) signal both around the antennal lobes and within them (Fig. 3A). These results mirror the time course of Mmp-1 protein upregulation following antennal and maxillary palp injury, which is robust 1-day post-injury (dpi)(22). Moreover, labeling of ensheathing glial cell membranes with GFP allows for the localization of *mmp-1* mRNAs and associated ribosomes within the antennal lobes 20hpi. To quantify these results, a total fluorescence intensity analysis of each signal was completed within the antennal lobes. Spot detection was not possible, as many transcripts and ribosomes were located close to, or even overlapping, each other. The total intensity of both *mmp-1* transcript and ribosome-association signals significantly increased within antennal lobes 20hpi (Fig. 3B).

**Figure 3:**
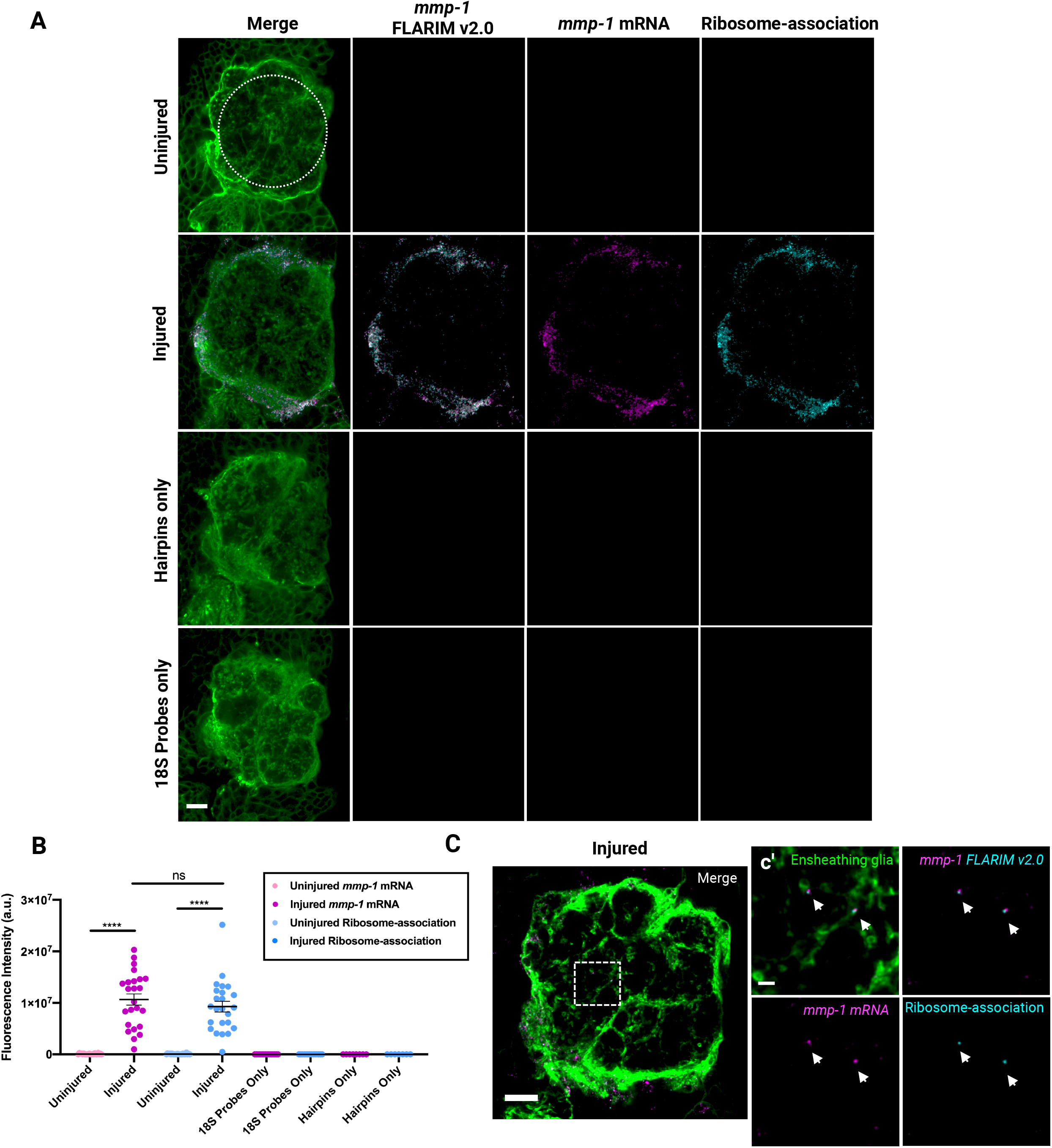
FLARIM v2.0 to detect *mmp-1* in the *Drosophila* ORN injury model. **(A)** Representative images of *mmp-1* FLARIM v2.0 probeset in uninjured and injured ALs with ROIs quantified (dotted circle). Injured: 20hpi. Hairpins only: 20hpi (no probes, only amplification with fluorescent hairpins). 18S Probes only: 20hpi (18S FLARIM probes with one half of the split initiator sequence and amplification with fluorescent hairpins). Single slice: 0.5µm. Scale bar: 20µm. Genotype: *;UAS-mCD8::GFP;TIFR-Gal4*. **(B)** Quantification of *mmp-1* FLARIM v2.0 total intensity (a.u.) in ALs. *mmp-1* mRNA Uninjured (n=22), *mmp-1* mRNA Injured (n=24), Ribosome-association Uninjured (n=22), Ribosome-association Injured (n=24), *mmp-1* mRNA 18S Probes Only (n=16), Ribosome-association 18S Probes Only (n=14), *mmp-1* mRNA Hairpins Only (n=8), Ribosome-association Hairpins Only (n=7). Mean ± SEM; Uninjured *mmp-1* mRNA and Injured *mmp-1* mRNA: t-test ****(p<0.0001); Uninjured Ribosome-association and Injured Ribosome-association: Mann-Whitney test ****(p<0.0001); non-significant (ns). **(C)** Representative image of *mmp-1* FLARIM v2.0 probeset in an injured AL (20hpi). Single slice: 0.5µm. Scale bar: 10µm. Genotype: *;UAS-mCD8::GFP;TIFR-Gal4*. (c’) *mmp-1* transcripts are associated with ribosomes in ensheathing glial processes (white arrows). Single slice: 0.5µm. Scale bar: 2µm.

A series of control experiments were also conducted to ascertain the specificity of the probesets and our signal. In the absence of hybridized probes, the hairpins amplified alone do not generate a fluorescent signal (Fig. 3A, B). Additionally, hybridization of only the 18S FLARIM probes does not result in a signal, as only half of the initiator sequence is present, and hairpins are unable to open and generate a fluorescent signal (Fig. 3A, B). We also determined that the *mmp-1* probeset was specific to our transcript of interest. Upon Mmp-1 RNAi knockdown in glial cells, *mmp-1* mRNA and ribosome-association signals were significantly diminished in the injured condition (Fig. 4A, B, C).

**Figure 4:**
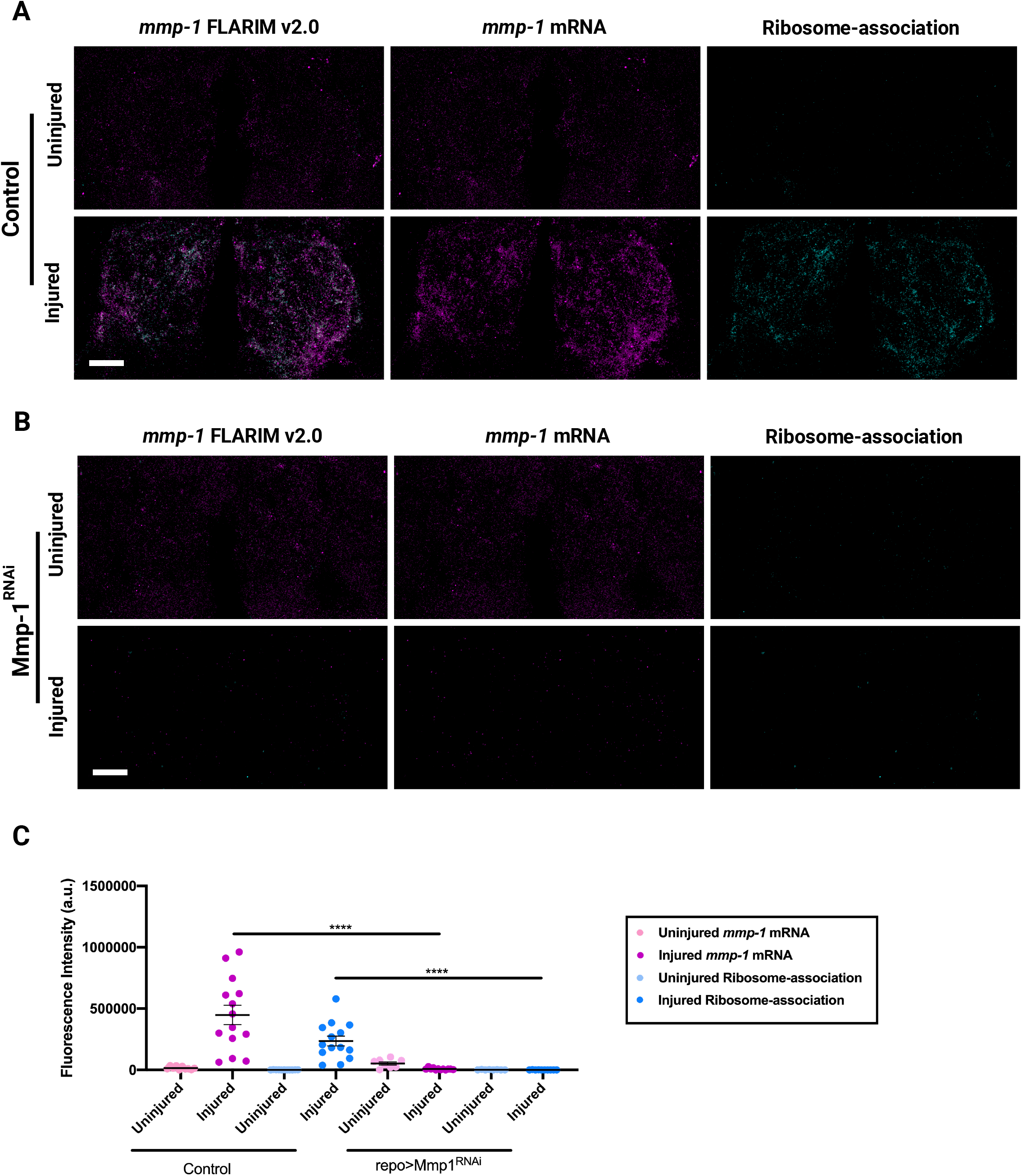
FLARIM v2.0 is specific to the *mmp-1* transcript and its associated ribosomes. **(A)** Representative images of *mmp-1* FLARIM v2.0 in control uninjured and injured (20hpi) brains. MIP: 15.5µm. Scale bar: 20µm. Genotype: *;tub-Gal80*^*ts*^*/+;repo-Gal4/+*. **(B)** Representative images of *mmp-1* FLARIM v2.0 in brains expressing Mmp-1 RNAi in allglial cells (repo>Mmp-1^RNAi^). MIP: 15.5µm. Scale bar: 20µm. Genotype: *;tub-Gal80*^*ts*^*/UAS-Mmp-1*^*RNAi*^*;repo-Gal4/+*. **(C)** Quantification of *mmp-1* FLARIM v2.0 total intensity (a.u.) in ALs. Control: Uninjured *mmp-1* mRNA (n=10), Injured *mmp-1* mRNA (n=14), Uninjured Ribosome-association (n=11), Injured Ribosome-association (n=14). repo>Mmp-1^RNAi^: Uninjured *mmp-1* mRNA (n=8), Injured *mmp-1* mRNA (n=12), Uninjured Ribosome-association (n=8), Injured Ribosome-association (n=10). Mean ± SEM; Mann-Whitney test ****(p<0.0001).

### Translation machinery is present in ensheathing glial processes

We observe that *mmp-1* transcripts and associated ribosomes are localized to ensheathing glial processes following ORN injury. As a result, we hypothesize that transcript localization and local translation of immune genes may be mechanisms employed by glial cells to mediate responses after a neuronal injury event. This is of particularly interest in relation to Mmp-1, as it is a secreted molecule employed by ensheathing glia to remodel the surrounding extracellular matrix and provide access to sites of damage(22, 23). However, in order for this to occur, translation and secretory machinery must also be present in ensheathing glial processes to properly modify and secrete Mmp-1(27). Evidence for translation machinery in glial processes has been demonstrated by previous groups. A study in mouse astrocytes has shown that along with a local pool of transcripts, endoplasmic reticulum (ER) and Golgi apparatus (GA) are also present in perivascular processes and endfeet(27). Therefore, we performed immunohistochemistry in uninjured and injured brains to assess whether the aforementioned organelles are present in glial projections of the ALs. Super-resolution microscopy allowed for the visualization of ER and GA within ensheathing glial fine processes (Fig. 5A-D). We were able to observe that ER and GA are present under both basal conditions and following ORN injury (Fig. 5A-D). These results further suggest that transcript localization and local translation of immune genes could be supported in glial processes, and could play a functional role in mediating rapid glial responses following neuronal injury

**Figure 5:**
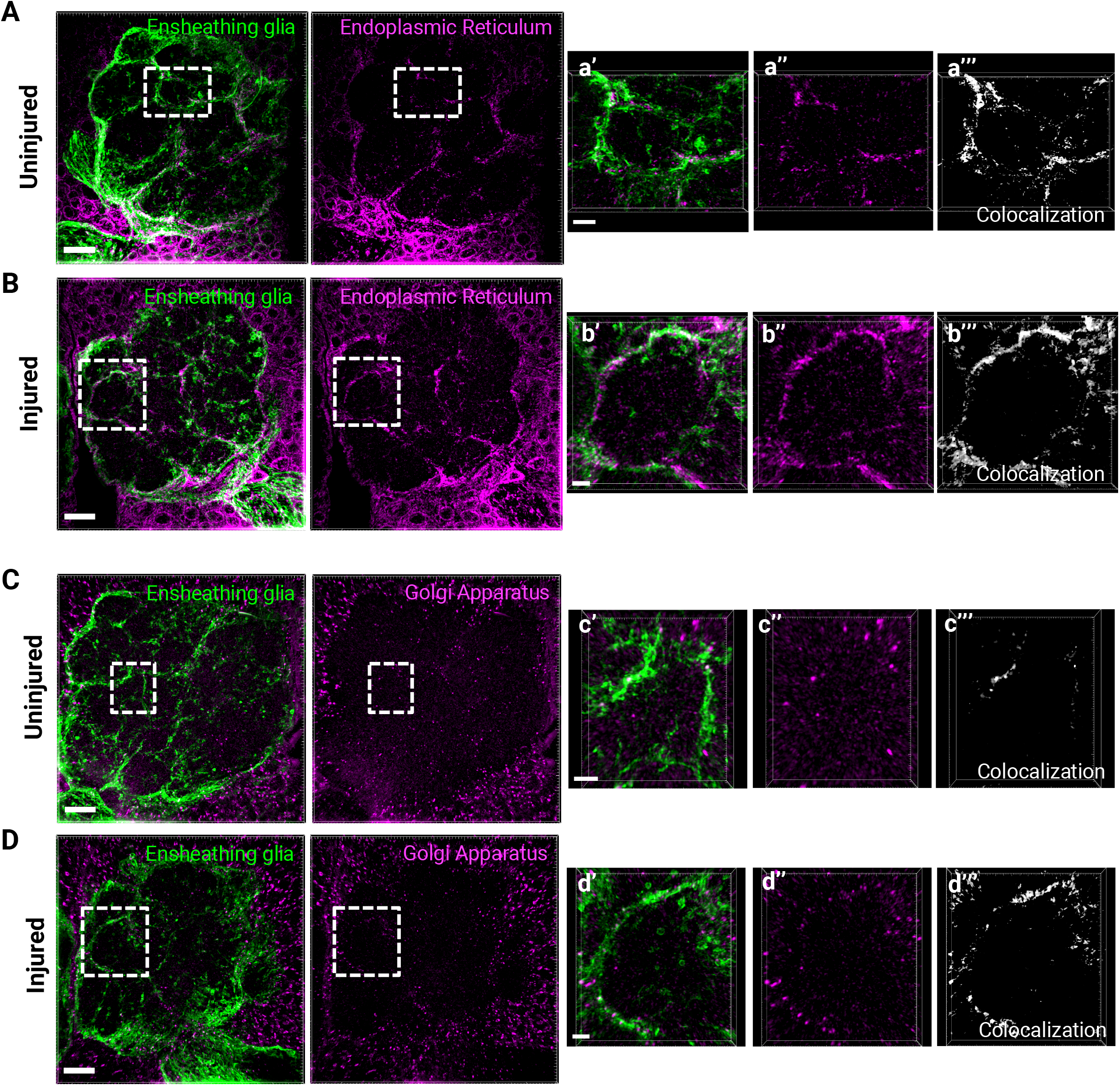
Endoplasmic reticulum and Golgi apparatus are present in ensheathing glial fine processes. **(A)** Representative super-resolution images of ER immunostaining in an uninjured AL. MIP (1.75µm). Scale bar: 10µm. Genotype: *;UAS-mCD8::GFP/UAS-mCD8::GFP;TIFR-Gal4/TIFR-Gal4*. **(a’, a’’)** ER is present within ensheathing glial fine processes of an uninjured AL. MIP (1.75µm). Scale bar: 3µm. **(a’’’)** Colocalization of ensheathing glia and ER signal. **(B)** Representative super-resolution images of ER immunostaining in an injured (20hpi) AL. MIP (1.75µm). Scale bar: 10µm. Genotype: *;UAS-mCD8::GFP/UAS-mCD8::GFP;TIFR-Gal4/TIFR-Gal4*. **(b’, b’’)** ER is present within ensheathing glial fine processes of an injured AL. MIP (1.75µm). Scale bar: 2µm. **(b’’’)** Colocalization of ensheathing glia and ER signal. **(C)** Representative super-resolution images of GA immunostaining in an uninjured AL. MIP (1.75µm). Scale bar: 10µm. Genotype: *;UAS-mCD8::GFP/UAS-mCD8::GFP;TIFR-Gal4/TIFR-Gal4*. **(c’, c’’)** GA is present within ensheathing glial fine processes of an uninjured AL. MIP (1.75µm). Scale bar: 2µm. **(c’’’)** Colocalization of ensheathing glia and GA signal. **(D)** Representative super-resolution images of GA immunostaining in an injured (20hpi) AL. MIP (1.75µm). Scale bar: 10µm. Genotype: *;UAS-mCD8::GFP/UAS-mCD8::GFP;TIFR-Gal4/TIFR-Gal4*. **(d’, d’’)** GA is present within ensheathing glial fine processes of an injured AL. MIP (1.75µm). Scale bar: 2µm. **(d’’’)** Colocalization of ensheathing glia and GA signal.

## Discussion

In this study, we have modified HCR and FLARIM methods to develop a novel dual probeset strategy to visualize single mRNA transcripts and their associated ribosomes *in vivo*(30, 39, 41). Specifically, we have applied this approach to visualize Mmp-1 transcripts in ensheathing glial cells responding to acute axon injury in adult *Drosophila*. It is becoming increasingly clear that directed transport of select mRNAs and local translation are essential for a wide range of cellular responses, including glial cell function in both the developing and mature brain(54-57). Thus, we propose that this novel methodology to visualize and quantify transcripts and, notably, ribosome association in whole tissue will be broadly valuable.

The mechanisms of mRNA localization and local translation are highly conserved and provide cells with the ability to restrict responses to subcellular compartments, conserve energy, and localize responses to specific stimuli[49]. Directed transport/translation of transcripts has been well described in neurons(58-62). More recently, the importance of local translation has been explored in glial cells. Local transcriptomes and translatomes have been characterized within astrocyte peripheral processes at the tripartite synapse, as well as astrocytic endfeet at the gliovascular interface(27, 28). In oligodendrocytes, myelin basic protein (*mbp*) mRNA is localized to oligodendrocyte sheaths and is required for proper myelin and axon development, while radial glia transport and locally translate mRNAs in their distal processes during development(25, 55). Research has also focused on how specific stimuli activate such mechanisms. Peripheral processes in astrocytes induce local translation in response to fear conditioning in mice, while neuronal activity may control myelination by oligodendrocytes through the induction of local protein synthesis(26, 29).

Here, we show that that our *mmp-1* FLARIM v2.0 approach can reliably detect significant *mmp-1* upregulation in glia following olfactory nerve injury and that this strategy allows us to monitor changes in the spatial distribution of mmp-1 transcripts, as well as association with ribosomes within one day after axotomy. We propose that our detection of transcript/ribosome association indicates that local translation of secreted Mmp-1 protein occurs at the distal process of glial cells as they invade injury sites (Fig. 3). We observe markers for both ER and GA in fine distal processes of ensheathing glia under both basal and injury conditions, suggesting that this glial subtype is equipped with organelles to locally translate and secrete a released factor such as Mmp-1 (Fig. 5). Given the ramified morphology of ensheathing glia, and the well-known role of Mmp-1 in ECM remodeling, rapid, local production of Mmp-1 is likely important for proper extension of glial processes within the deeper regions of the antennal lobes. As a result, this would aid phagocytic engulfment of degenerating olfactory neurons.

From a technical standpoint, FLARIM v2.0 offers a number of important features and, notably, some key advantages over the original FLARIM approach (Fig. 2). Fewer probes are required to monitor association of ribosomes and transcripts of interest. Transcripts and ribosomes are visualized as diffraction-limited spots and can be quantified to assess relative changes in gene expression and ribosome association in discrete cell types. Moreover, this method allows for flexible fluorescent labeling within samples. Finally, the generation of probes ordered as oPools is very cost effective. This and related protocols also offer the obvious benefit of greater spatial resolution to quantify gene upregulation in discrete cell types in heterogenous tissue, as opposed to cruder approaches (e.g., quantitative-PCR on crushed tissue).

More broadly, this approach offers a novel experimental readout for a component of glial immunity that has not yet been explored in flies and minimally in other model organisms, namely local translation of immune genes at injury sites. Our results now offer a new platform to explore future questions to investigate how mRNA transcripts are transported, translated, and eventually degraded once an immune response is no longer required. Previous work from our lab has shown that insulin-like signaling (ILS) pathways are acutely activated in ensheathing glial cells responding to olfactory nerve axotomy(63). Because ILS cascades are known to promote protein translation via mammalian target of rapamycin (mTOR), this offers an intriguing candidate by which immune gene translation is locally enhanced within glia in response to neurodegeneration(64). Future screening efforts will reveal how select immune gene transcripts are shuttled throughout glial processes to support proper innate immune reactions to acute trauma and perhaps also chronic neurodegenerative conditions.

## Methods

### Probe design and synthesis

Probes to the transcript of interest were designed using Oligostan in R Studio(40, 65). The probes were compared to a database of noncoding RNAs (http://flybase.org(66): http://ftp.flybase.net/genomes/Drosophila_melanogaster/dmel_r6.37_FB2020_06/fasta/; dmel-all-miRNA-r6.37.fasta.gz, dmel-all-miscRNA-r6.37.fasta.gz, dmel-all-ncRNA-r6.37.fasta.gz) for sequence similarity using Geneious (Biomatters). Any probes similar to ncRNAs (miRNA, snoRNA, lncRNA) were discarded from the probeset. The remaining probes were used to make up the gene-specific probesets. B2 and split B3 initiator sequences were added to the probes to allow for HCR amplification and FLARIM detection, which have been previously described(30, 39). Probes were synthesized in 96-well plates or as 50 pmol oPools (Integrated DNA Technologies). Probe sequences can be found in Supplementary tables 1-3.

### ORN Injury Assay and Dissection

Third antennal segments and maxillary palps were removed bilaterally using forceps, as previously described(45). Flies were raised at 25°C and returned to this temperature following injury, until dissection at 20hpi. Fly lines containing the tubulin-Gal80ts transgene were raised at 18°C and moved to 30°C for 7 days to induce Mmp-1 RNAi expression. These flies were then injured and moved back to 30°C until dissection. Fly heads were pulled and fixed in 4% paraformaldehyde (PFA) + 0.1% Triton X-100 for 20min at room temperature (RT) on a rocker. Then, heads were washed in 1X PBS-TX (0.01% Triton X-100) on a rocker at RT (3×2min). Brains were dissected in 1X PBS-TX (0.01% Triton X-100). Brains were then fixed in 4% PFA + 0.1% Triton X-100 for 20min at RT on a rocker and washed with 1X PBS-TX (0.1% Triton X-100) on a rocker at RT (3×2min). To increase probe penetration, brains were also permeabilized for 20min in 1X PBS-TX (0.5% Triton X-100), while rocking at RT. Brains for immunostaining were not additionally permeabilized.

### smiFISH

The smiFISH reagents, flap hybridization, and protocol have been described previously(37, 40). The smiFISH protocol has been adapted as follows: After dissection, fixation, and permeabilization, brains were placed in hybridization buffer(40) with the Mmp-1 smiFISH ATTO 550 (Integrated DNA Technologies) probeset and hybridized overnight in a thermal cycler at 37°C. The following day, brains were washed in 1X SSC/10% formamide/0.1% Triton X-100/0.1% Tween-20 at 50°C (2×30min) in a thermal cycler. Brains were then mounted in Vectashield (Vector Labs) under #1.5 coverslips and imaged.

### Probe Hybridization and Signal Amplification

The HCR v3.0 hybridization and amplification protocol has been described previously for whole-mount fruit fly embryos(41, 67). The protocol was utilized in adult fruit fly brains as follows: After permeabilization, brains were pre-hybridized for 20min at 37°C in probe hybridization buffer (Molecular Instruments), which had already been heated to 37°C. Brains were then hybridized overnight (16-18hrs) with the gene-specific and 18S FLARIM probes in a thermal cycler at 37°C. The following day, brains were washed in probe wash buffer (Molecular Instruments) at 37°C in the thermal cycler (4×15min). During the washes, HCR hairpins (Molecular Instruments) were snap-cooled, and the amplification buffer (Molecular Instruments) was moved to RT. Brains were washed again in 5X SSCT (0.1% Tween-20) at RT (2×5min). Brains were pre-amplified with amplification buffer at RT for 10min. Then, hairpins were added to the amplification buffer and the HCR and FLARIM signals were amplified. Amplification times were empirically determined for each probeset in this paper (*mmp-1* HCR B2 FLARIM split B3 Alexa Fluor 546: 1hr; 18S FLARIM B3 Alexa Fluor 647: 6hrs). Following amplification, brains were washed in 5X SSCT (0.1% Tween-20) at RT (5min, 2×30min, 5min), and then mounted in Vectashield (Vector Labs) under #1.5 coverslips and imaged.

### Immunostaining

Antibodies were diluted in 1X PBS-TX (0.1% Triton X-100) and brains were incubated in primary antibodies overnight on a shaker at 4°C. Brains were washed in PBS-TX (0.1% Triton X-100) at RT (3×30min) and incubated in secondary antibodies in 1X PBS-TX (0.1% Triton X-100) for 2hrs at RT on a shaker. Brains were then washed again in 1X PBS-TX (0.1% Triton X-100) at RT (3×30min) and mounted in Vectashield (Vector Labs) under #1.5 coverslips and imaged.

### Antibodies

The following primary antibodies were used: mouse beta-galactosidase 40-1a (Developmental Studies Hybridoma Bank) at 1:100, chicken anti-GFP (ThermoFisher, #A10262) at 1:1000, mouse anti-nc82 (Bruchpilot; Developmental Studies Hybridoma Bank) at 1:50, goat anti-GMAP (Developmental Studies Hybridoma Bank) at 1:800, mouse anti-Cnx99A 6-2-1 (Developmental Studies Hybridoma Bank) at 1:400. All secondary antibodies (Jackson Immunoresearch 703-545-155, 715-295-150, and 705-295-147) were used at a 1:400 dilution.

### Microscopy and Analysis

Samples were imaged on a Zeiss LSM 700 with a Zeiss 40X 1.4 NA oil immersion plan-apochromatic lens. Samples with Golgi and ER staining were imaged using a Zeiss Elyra 7 with lattice SIM with a Zeiss 63X 1.4 NA oil immersion plan-apochromatic lens at the OHSU Advanced Light Microscopy Core (ALMC). Brains within the same experiment were imaged on the same day, using the same microscope settings. smiFISH and FLARIM v2.0 images were pre-processed by smooth filtering in Zen, while Golgi and ER images were SIM processed in Zen. Volocity 3D Image Analysis Software (Quorum Technologies) was used for fluorescence quantification, while Imaris Cell Imaging Software (Andor Technology) at the OHSU ALMC was used for super-resolution image processing and colocalization analysis. GraphPad Prism 8 (Graphpad Software) was used for statistical analysis: Student’s t-test, Mann-Whitney test. Normality was tested using the D’Agostino-Pearson normality test. Outliers were identified using the ROUT method. Experiments were not blinded. For *mmp-1* FLARIM v2.0 quantifications, 5um was removed from the top of each z-stack to exclude ensheathing glial cell bodies from the analysis, and only quantify *mmp-1* mRNA and ribosome-association signals within the ensheathing glial processes of the ALs. A circular ROI was defined in each AL, where the total fluorescence for each signal was calculated.

### Drosophila Stocks

Adult flies between 4-14 days old were used for experiments. The following lines were used: w1118 (BDSC 5905), UAS-mCD8::GFP (BDSC 5137), UAS-LacZ::NLS (BDSC 3956), TIFR-Gal4(68), repo-Gal4(12), tubulin-Gal80ts (BDSC 7108), UAS-Mmp-1 RNAi(69).

## Acknowledgements

We acknowledge expert technical assistance by staff in the Advanced Light Microscopy Core, supported by grant P30NS061800. We thank Alexandra Houser for help with smiFISH experiments. We would also like to thank Dirk Bohmann, Bloomington Drosophila Stock Center at Indiana University, and the Developmental Studies Hybridoma Bank at the University of Iowa for flies and antibodies.

## Funding

This work was supported by National Institutes of Health grants (R21 NS084112, R21 NS107771, and R01 NS117934), the Ken and Ginger Term Professor Award (M.A.L.), and the Tartar Trust Fellowship at Oregon Health and Sciences University (P.R.).

## Conflict of Interest

The authors declare that no competing interests exist.

---

**Supplementary Figure 1:**
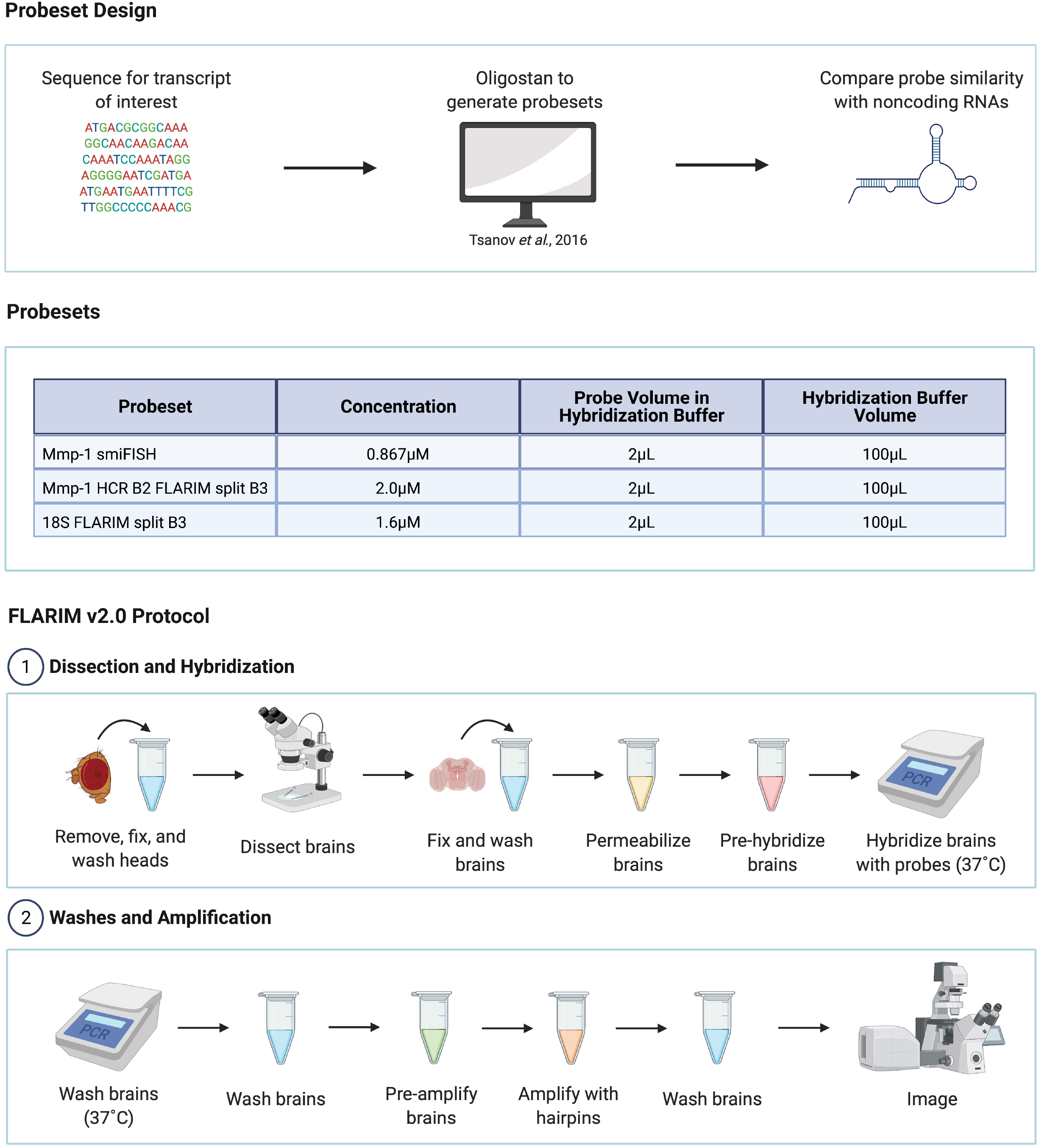
Probe design and protocol workflow. <u>Probeset design</u>: Probesets to the transcript of interest were designed using Oligostan (Tsanov *et al*., 2016) and sequence similarity was compared to noncoding RNAs. <u>Probesets</u>: Probeset sequences are listed in in Supplementary tables 1, 2, and 3. The smiFISH probeset assembly, protocol, and reagents are described in Tsanov *et al*., 2016. For the Mmp-1 HCR B2 FLARIM split B3 and 18S FLARIM split B3 probesets, the FLARIM v2.0 protocol was used (*see Methods for details*) <u>FLARIM v2.0 Protocol</u>: Dissection and hybridization: fly brains are dissected and hybridized overnight with gene-specific and 18S ribosome probes in a thermal cycler. Washes and amplification: the following day, brains are washed and amplified with fluorescent hairpins to generate mRNA-specific and ribosome-association signals (*see Methods for details*). Created with BioRender.com (Adapted from “Growing Norovirus in Human Intestinal Enteroids (HIEs)”, by BioRender.com (2020). Retrieved from https://app.biorender.com/biorender-templates)

**Supplementary Table 1:**
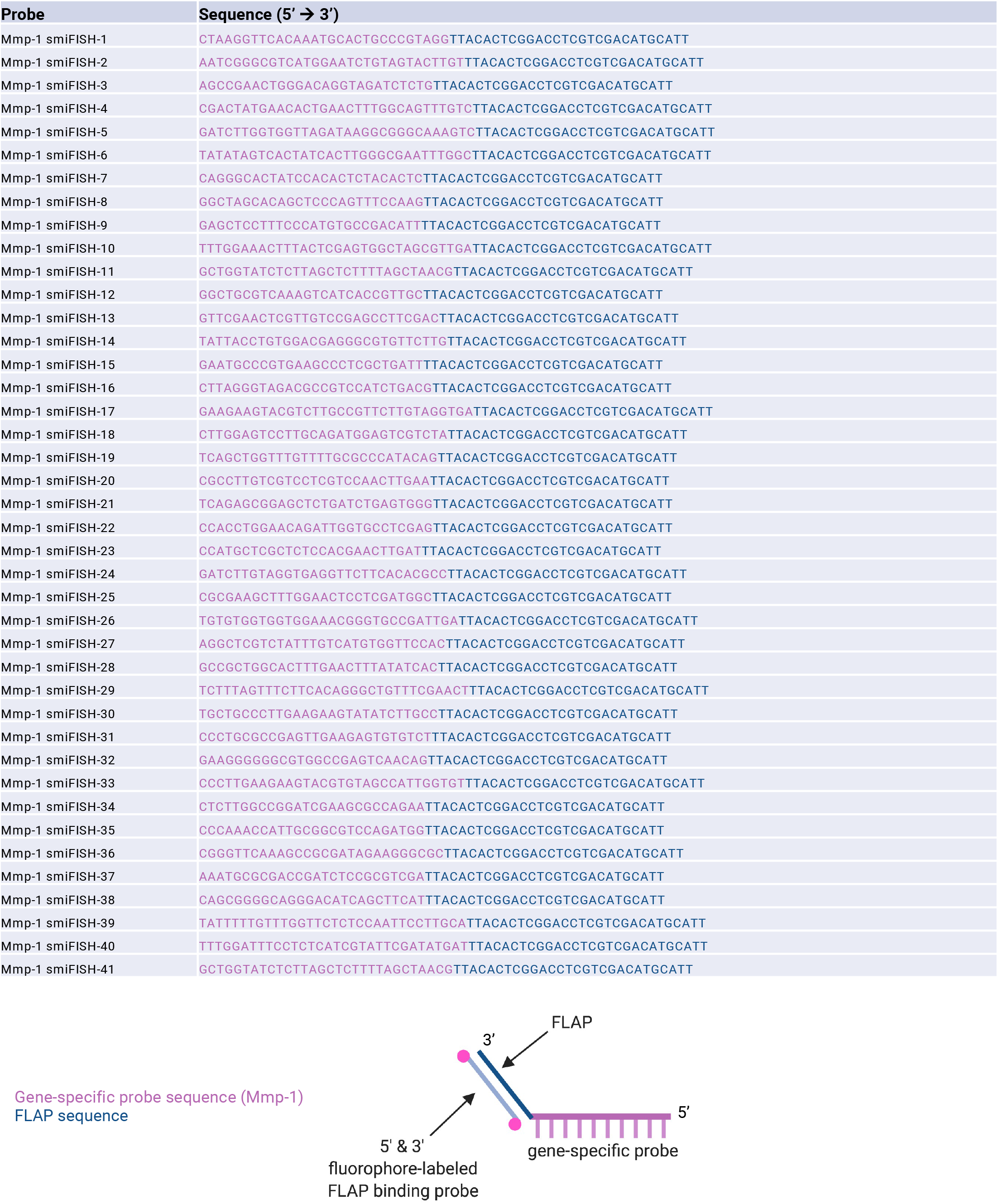
smiFISH *mmp-1* probe sequences. *mmp-1* gene-specific sequences (magenta), FLAP sequence (blue; Tsanov, *et al*., 2016)

**Supplementary Table 2:**
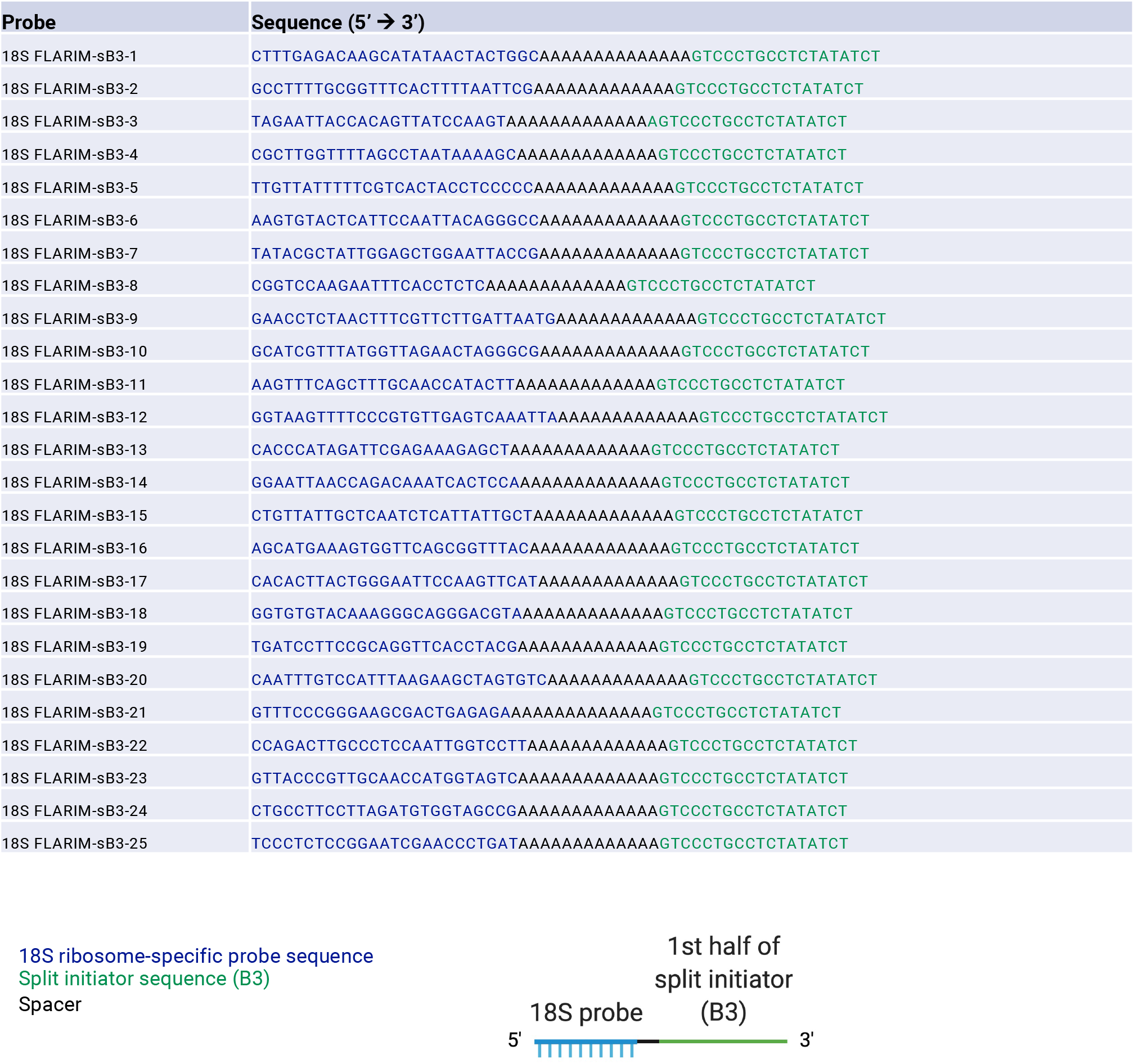
18S FLARIM split B3 probe sequences. 18S ribosome-specific probe sequences (blue) and B3 split initiator sequence (green; Choi *et al*., 2014).

**Supplementary Table 3:**
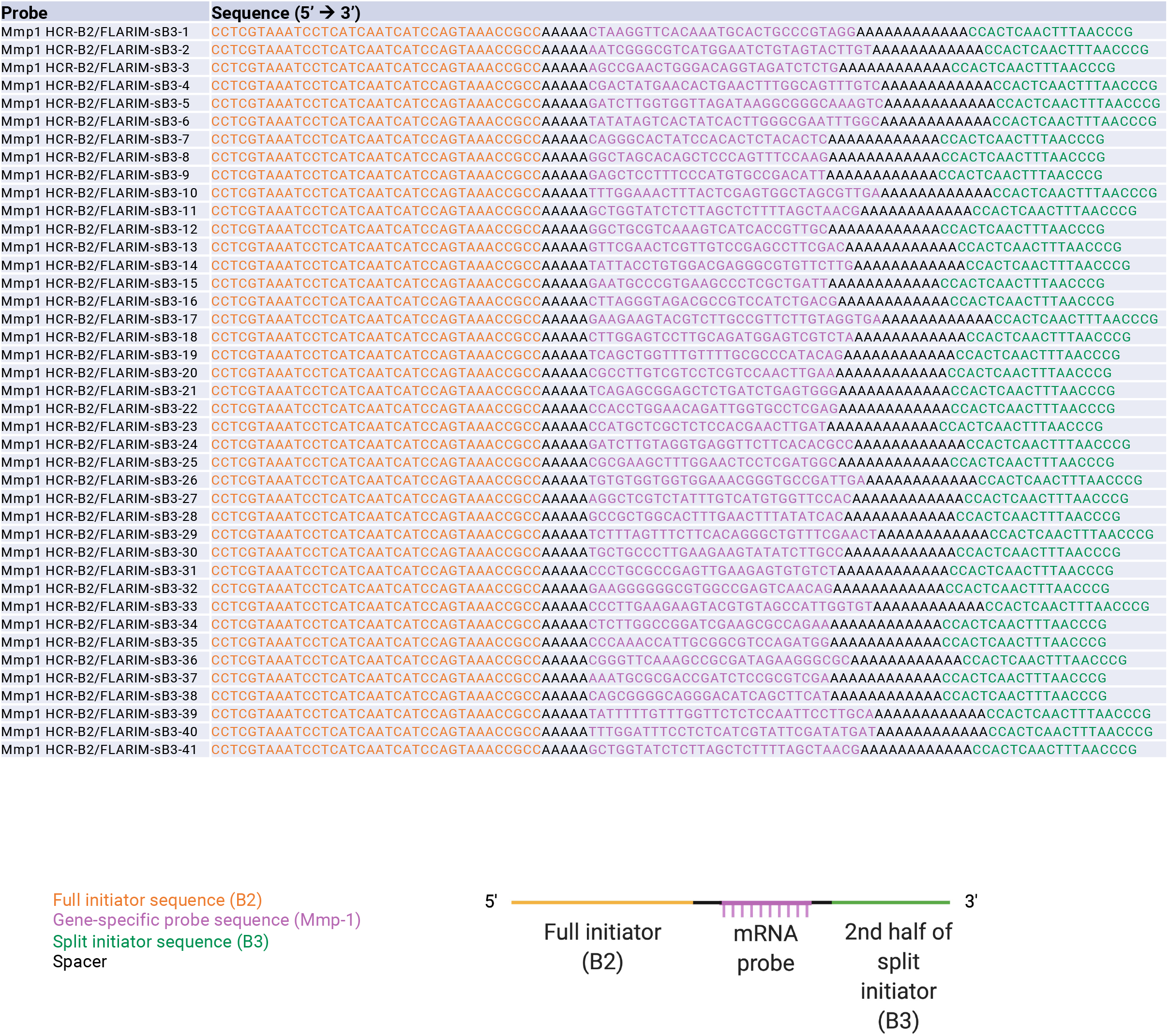
*mmp-1* HCR B2 FLARIM split B3 probe sequences. *mmp-1* gene-specific sequences (magenta), B3 split initiator sequence (green; Choi *et al*., 2014), and full B2 initiator sequence (orange; Choi *et al*., 2014).

## Notes

### Competing Interest Statement

The authors have declared no competing interest.

## References

1. Fields RD, Stevens-Graham B. New insights into neuron-glia communication. Science. 2002;298(5593):556–62. 10.1126/science.298.5593.556

2. Barres BA. The mystery and magic of glia: a perspective on their roles in health and disease. Neuron. 2008;60(3):430–40. 10.1016/j.neuron.2008.10.013

3. Allen NJ, Barres BA. Neuroscience: Glia - more than just brain glue. Nature. 2009;457(7230):675–7. 10.1038/457675a

4. Napoli I, Neumann H. Microglial clearance function in health and disease. Neuroscience. 2009;158(3):1030–8. 10.1016/j.neuroscience.2008.06.046

5. Polazzi E, Monti B. Microglia and neuroprotection: from in vitro studies to therapeutic applications. Prog Neurobiol. 2010;92(3):293–315. 10.1016/j.pneurobio.2010.06.009

6. Hong S, Stevens B. Microglia: Phagocytosing to Clear, Sculpt, and Eliminate. Dev Cell. 2016;38(2):126–8. 10.1016/j.devcel.2016.07.006

7. Glezer I, Simard AR, Rivest S. Neuroprotective role of the innate immune system by microglia. Neuroscience. 2007;147(4):867–83. 10.1016/j.neuroscience.2007.02.055

8. Ray A, Speese SD, Logan MA. Glial Draper Rescues Abeta Toxicity in a Drosophila Model of Alzheimer’s Disease. J Neurosci. 2017;37(49):11881–93. 10.1523/JNEUROSCI.0862-17.2017

9. Freeman MR, Doherty J. Glial cell biology in Drosophila and vertebrates. Trends Neurosci. 2006;29(2):82–90. 10.1016/j.tins.2005.12.002

10. Ziegenfuss JS, Doherty J, Freeman MR. Distinct molecular pathways mediate glial activation and engulfment of axonal debris after axotomy. Nat Neurosci. 2012;15(7):979–87. 10.1038/nn.3135

11. Logan MA. Glial contributions to neuronal health and disease: new insights from Drosophila. Curr Opin Neurobiol. 2017;47:162–7. 10.1016/j.conb.2017.10.008

12. MacDonald JM, Beach MG, Porpiglia E, Sheehan AE, Watts RJ, Freeman MR. The Drosophila cell corpse engulfment receptor Draper mediates glial clearance of severed axons. Neuron. 2006;50(6):869–81. 10.1016/j.neuron.2006.04.028

13. Mishra B, Carson R, Hume RI, Collins CA. Sodium and potassium currents influence Wallerian degeneration of injured Drosophila axons. J Neurosci. 2013;33(48):18728–39. 10.1523/JNEUROSCI.1007-13.2013

14. Freeman MR. Signaling mechanisms regulating Wallerian degeneration. Curr Opin Neurobiol. 2014;27:224–31. 10.1016/j.conb.2014.05.001

15. Awasaki T, Tatsumi R, Takahashi K, Arai K, Nakanishi Y, Ueda R, Ito K. Essential role of the apoptotic cell engulfment genes draper and ced-6 in programmed axon pruning during Drosophila metamorphosis. Neuron. 2006;50(6):855–67. 10.1016/j.neuron.2006.04.027

16. Ziegenfuss JS, Biswas R, Avery MA, Hong K, Sheehan AE, Yeung YG, Stanley ER, Freeman MR. Draper-dependent glial phagocytic activity is mediated by Src and Syk family kinase signalling. Nature. 2008;453(7197):935–9. 10.1038/nature06901

17. Logan MA, Hackett R, Doherty J, Sheehan A, Speese SD, Freeman MR. Negative regulation of glial engulfment activity by Draper terminates glial responses to axon injury. Nat Neurosci. 2012;15(5):722–30. 10.1038/nn.3066

18. Scheib JL, Sullivan CS, Carter BD. Jedi-1 and MEGF10 signal engulfment of apoptotic neurons through the tyrosine kinase Syk. J Neurosci. 2012;32(38):13022–31. 10.1523/JNEUROSCI.6350-11.2012

19. Macdonald JM, Doherty J, Hackett R, Freeman MR. The c-Jun kinase signaling cascade promotes glial engulfment activity through activation of draper and phagocytic function. Cell Death Differ. 2013;20(9):1140–8. 10.1038/cdd.2013.30

20. Doherty J, Sheehan AE, Bradshaw R, Fox AN, Lu TY, Freeman MR. PI3K signaling and Stat92E converge to modulate glial responsiveness to axonal injury. PLoS Biol. 2014;12(11):e1001985. 10.1371/journal.pbio.1001985

21. Lu TY, MacDonald JM, Neukomm LJ, Sheehan AE, Bradshaw R, Logan MA, Freeman MR. Axon degeneration induces glial responses through Draper-TRAF4-JNK signalling. Nat Commun. 2017;8:14355. 10.1038/ncomms14355

22. Purice MD, Ray A, Munzel EJ, Pope BJ, Park DJ, Speese SD, Logan MA. A novel Drosophila injury model reveals severed axons are cleared through a Draper/MMP-1 signaling cascade. Elife. 2017;6. 10.7554/eLife.23611

23. Fujioka H, Dairyo Y, Yasunaga K, Emoto K. Neural functions of matrix metalloproteinases: plasticity, neurogenesis, and disease. Biochem Res Int. 2012;2012:789083. 10.1155/2012/789083

24. Doherty J, Logan MA, Tasdemir OE, Freeman MR. Ensheathing glia function as phagocytes in the adult Drosophila brain. J Neurosci. 2009;29(15):4768–81. 10.1523/JNEUROSCI.5951-08.2009

25. Lyons DA, Naylor SG, Scholze A, Talbot WS. Kif1b is essential for mRNA localization in oligodendrocytes and development of myelinated axons. Nat Genet. 2009;41(7):854–8. 10.1038/ng.376

26. Wake H, Lee PR, Fields RD. Control of local protein synthesis and initial events in myelination by action potentials. Science. 2011;333(6049):1647–51. 10.1126/science.1206998

27. Boulay AC, Saubamea B, Adam N, Chasseigneaux S, Mazare N, Gilbert A, Bahin M, Bastianelli L, Blugeon C, Perrin S, Pouch J, Ducos B, Le Crom S, Genovesio A, Chretien F, Decleves X, Laplanche JL, Cohen-Salmon M. Translation in astrocyte distal processes sets molecular heterogeneity at the gliovascular interface. Cell Discov. 2017;3:17005. 10.1038/celldisc.2017.5

28. Sakers K, Lake AM, Khazanchi R, Ouwenga R, Vasek MJ, Dani A, Dougherty JD. Astrocytes locally translate transcripts in their peripheral processes. Proc Natl Acad Sci U S A. 2017;114(19):E3830–E8. 10.1073/pnas.1617782114

29. Mazare N, Oudart M, Moulard J, Cheung G, Tortuyaux R, Mailly P, Mazaud D, Bemelmans AP, Boulay AC, Blugeon C, Jourdren L, Le Crom S, Rouach N, Cohen-Salmon M. Local Translation in Perisynaptic Astrocytic Processes Is Specific and Changes after Fear Conditioning. Cell Rep. 2020;32(8):108076. 10.1016/j.celrep.2020.108076

30. Burke KS, Antilla KA, Tirrell DA. A Fluorescence in Situ Hybridization Method To Quantify mRNA Translation by Visualizing Ribosome-mRNA Interactions in Single Cells. ACS Cent Sci. 2017;3(5):425–33. 10.1021/acscentsci.7b00048

31. Kosman D, Mizutani CM, Lemons D, Cox WG, McGinnis W, Bier E. Multiplex detection of RNA expression in Drosophila embryos. Science. 2004;305(5685):846. 10.1126/science.1099247

32. Pare A, Lemons D, Kosman D, Beaver W, Freund Y, McGinnis W. Visualization of individual Scr mRNAs during Drosophila embryogenesis yields evidence for transcriptional bursting. Curr Biol. 2009;19(23):2037–42. 10.1016/j.cub.2009.10.028

33. Weil TT, Parton RM, Herpers B, Soetaert J, Veenendaal T, Xanthakis D, Dobbie IM, Halstead JM, Hayashi R, Rabouille C, Davis I. Drosophila patterning is established by differential association of mRNAs with P bodies. Nat Cell Biol. 2012;14(12):1305–13. 10.1038/ncb2627

34. Abbaszadeh EK, Gavis ER. Fixed and live visualization of RNAs in Drosophila oocytes and embryos. Methods. 2016;98:34–41. 10.1016/j.ymeth.2016.01.018

35. Long X, Colonell J, Wong AM, Singer RH, Lionnet T. Quantitative mRNA imaging throughout the entire Drosophila brain. Nat Methods. 2017;14(7):703–6. 10.1038/nmeth.4309

36. Trcek T, Lionnet T, Shroff H, Lehmann R. mRNA quantification using single-molecule FISH in Drosophila embryos. Nat Protoc. 2017;12(7):1326–48. 10.1038/nprot.2017.030

37. Yang L, Titlow J, Ennis D, Smith C, Mitchell J, Young FL, Waddell S, Ish-Horowicz D, Davis Single molecule fluorescence in situ hybridisation for quantitating post-transcriptional regulation in Drosophila brains. Methods. 2017;126:166–76. 10.1016/j.ymeth.2017.06.025

38. Meissner GW, Nern A, Singer RH, Wong AM, Malkesman O, Long X. Mapping Neurotransmitter Identity in the Whole-Mount Drosophila Brain Using Multiplex High-Throughput Fluorescence in Situ Hybridization. Genetics. 2019;211(2):473–82. 10.1534/genetics.118.301749

39. Choi HM, Beck VA, Pierce NA. Multiplexed in situ hybridization using hybridization chain reaction. Zebrafish. 2014;11(5):488–9. 10.1089/zeb.2014.1501

40. Tsanov N, Samacoits A, Chouaib R, Traboulsi AM, Gostan T, Weber C, Zimmer C, Zibara K, Walter T, Peter M, Bertrand E, Mueller F. smiFISH and FISH-quant - a flexible single RNA detection approach with super-resolution capability. Nucleic Acids Res. 2016;44(22):e165. 10.1093/nar/gkw784

41. Choi HMT, Schwarzkopf M, Fornace ME, Acharya A, Artavanis G, Stegmaier J, Cunha A, Pierce NA. Third-generation in situ hybridization chain reaction: multiplexed, quantitative, sensitive, versatile, robust. Development. 2018;145(12). 10.1242/dev.165753

42. Ayaz D, Leyssen M, Koch M, Yan J, Srahna M, Sheeba V, Fogle KJ, Holmes TC, Hassan BA. Axonal injury and regeneration in the adult brain of Drosophila. J Neurosci. 2008;28(23):6010–21. 10.1523/JNEUROSCI.0101-08.2008

43. Vosshall LB, Amrein H, Morozov PS, Rzhetsky A, Axel R. A spatial map of olfactory receptor expression in the Drosophila antenna. Cell. 1999;96(5):725–36.

44. Logan MA, Freeman MR. The scoop on the fly brain: glial engulfment functions in Drosophila. Neuron Glia Biol. 2007;3(1):63–74. 10.1017/S1740925X07000646

45. Purice MD, Speese SD, Logan MA. Delayed glial clearance of degenerating axons in aged Drosophila is due to reduced PI3K/Draper activity. Nat Commun. 2016;7:12871. 10.1038/ncomms12871

46. Freeman MR. Drosophila Central Nervous System Glia. Cold Spring Harb Perspect Biol. 2015;7(11). 10.1101/cshperspect.a020552

47. Kremer MC, Jung C, Batelli S, Rubin GM, Gaul U. The glia of the adult Drosophila nervous system. Glia. 2017;65(4):606–38. 10.1002/glia.23115

48. Wu B, Li J, Chou YH, Luginbuhl D, Luo L. Fibroblast growth factor signaling instructs ensheathing glia wrapping of Drosophila olfactory glomeruli. Proc Natl Acad Sci U S A. 2017;114(29):7505–12. 10.1073/pnas.1706533114

49. Dziembowska M, Wlodarczyk J. MMP9: a novel function in synaptic plasticity. Int J Biochem Cell Biol. 2012;44(5):709–13. 10.1016/j.biocel.2012.01.023

50. Dziembowska M, Milek J, Janusz A, Rejmak E, Romanowska E, Gorkiewicz T, Tiron A, Bramham CR, Kaczmarek L. Activity-dependent local translation of matrix metalloproteinase-9. J Neurosci. 2012;32(42):14538–47. 10.1523/JNEUROSCI.6028-11.2012

51. Tantale K, Mueller F, Kozulic-Pirher A, Lesne A, Victor JM, Robert MC, Capozi S, Chouaib R, Backer V, Mateos-Langerak J, Darzacq X, Zimmer C, Basyuk E, Bertrand E. A single-molecule view of transcription reveals convoys of RNA polymerases and multi-scale bursting. Nat Commun. 2016;7:12248. 10.1038/ncomms12248

52. Dermit M, Dodel M, Mardakheh FK. Methods for monitoring and measurement of protein translation in time and space. Mol Biosyst. 2017;13(12):2477–88. 10.1039/c7mb00476a

53. Biswas J, Liu Y, Singer RH, Wu B. Fluorescence Imaging Methods to Investigate Translation in Single Cells. Cold Spring Harb Perspect Biol. 2019;11(4). 10.1101/cshperspect.a032722

54. Martin KC, Ephrussi A. mRNA localization: gene expression in the spatial dimension. Cell. 2009;136(4):719–30. 10.1016/j.cell.2009.01.044

55. Pilaz LJ, Lennox AL, Rouanet JP, Silver DL. Dynamic mRNA Transport and Local Translation in Radial Glial Progenitors of the Developing Brain. Curr Biol. 2016;26(24):3383–92. 10.1016/j.cub.2016.10.040

56. Blanco-Urrejola M, Gaminde-Blasco A, Gamarra M, de la Cruz A, Vecino E, Alberdi E, Baleriola J. RNA Localization and Local Translation in Glia in Neurological and Neurodegenerative Diseases: Lessons from Neurons. Cells. 2021;10(3). 10.3390/cells10030632

57. Meservey LM, Topkar VV, Fu M-m. mRNA Transport and Local Translation in Glia. Trends in Cell Biology. 10.1016/j.tcb.2021.03.006

58. Akins MR, Berk-Rauch HE, Fallon JR. Presynaptic translation: stepping out of the postsynaptic shadow. Front Neural Circuits. 2009;3:17. 10.3389/neuro.04.017.2009

59. Kuklin EA, Alkins S, Bakthavachalu B, Genco MC, Sudhakaran I, Raghavan KV, Ramaswami M, Griffith LC. The Long 3’UTR mRNA of CaMKII Is Essential for Translation-Dependent Plasticity of Spontaneous Release in Drosophila melanogaster. J Neurosci. 2017;37(44):10554–66. 10.1523/JNEUROSCI.1313-17.2017

60. Rangaraju V, Tom Dieck S, Schuman EM. Local translation in neuronal compartments: how local is local? EMBO Rep. 2017;18(5):693–711. 10.15252/embr.201744045

61. Biever A, Donlin-Asp PG, Schuman EM. Local translation in neuronal processes. Curr Opin Neurobiol. 2019;57:141–8. 10.1016/j.conb.2019.02.008

62. Holt CE, Martin KC, Schuman EM. Local translation in neurons: visualization and function. Nat Struct Mol Biol. 2019;26(7):557–66. 10.1038/s41594-019-0263-5

63. Musashe DT, Purice MD, Speese SD, Doherty J, Logan MA. Insulin-like Signaling Promotes Glial Phagocytic Clearance of Degenerating Axons through Regulation of Draper. Cell Rep. 2016;16(7):1838–50. 10.1016/j.celrep.2016.07.022

64. Wang X, Proud CG. The mTOR pathway in the control of protein synthesis. Physiology (Bethesda). 2006;21:362–9. 10.1152/physiol.00024.2006

65. Team RS. R Studio: Integrated development for R. R Studio, PBC, Boston, MA 2020.

66. Thurmond J, Goodman JL, Strelets VB, Attrill H, Gramates LS, Marygold SJ, Matthews BB, Millburn G, Antonazzo G, Trovisco V, Kaufman TC, Calvi BR, FlyBase C. FlyBase 2.0: the next generation. Nucleic Acids Res. 2019;47(D1):D759–D65. 10.1093/nar/gky1003

67. Choi HM, Calvert CR, Husain N, Huss D, Barsi JC, Deverman BE, Hunter RC, Kato M, Lee SM, Abelin AC, Rosenthal AZ, Akbari OS, Li Y, Hay BA, Sternberg PW, Patterson PH, Davidson EH, Mazmanian SK, Prober DA, van de Rijn M, Leadbetter JR, Newman DK, Readhead C, Bronner ME, Wold B, Lansford R, Sauka-Spengler T, Fraser SE, Pierce NA. Mapping a multiplexed zoo of mRNA expression. Development. 2016;143(19):3632–7. 10.1242/dev.140137

68. Yao Y, Wu Y, Yin C, Ozawa R, Aigaki T, Wouda RR, Noordermeer JN, Fradkin LG, Hing H. Antagonistic roles of Wnt5 and the Drl receptor in patterning the Drosophila antennal lobe. Nat Neurosci. 2007;10(11):1423–32. 10.1038/nn1993

69. Uhlirova M, Bohmann D. JNK-and Fos-regulated Mmp1 expression cooperates with Ras to induce invasive tumors in Drosophila. EMBO J. 2006;25(22):5294–304. 10.1038/sj.emboj.7601401

